# Integrating animal movement with habitat suitability for estimating dynamic landscape connectivity

**DOI:** 10.1101/224766

**Authors:** Mariëlle L. van Toor, Bart Kranstauber, Scott H. Newman, Diann J. Prosser, John Y. Takekawa, Georgios Technitis, Robert Weibel, Martin Wikelski, Kamran Safi

## Abstract

**Context:** High-resolution animal movement data are becoming increasingly available, yet having a multitude of empirical trajectories alone does not allow us to easily predict animal movement. To answer ecological and evolutionary questions at a population level, quantitative estimates of a species’ potential to link patches or populations are of importance.

**Objectives:** We introduce an approach that combines movement-informed simulated trajectories with an environment-informed estimate of the trajectories’ plausibility to derive connectivity. Using the example of bar-headed geese we estimated migratory connectivity at a landscape level throughout the annual cycle in their native range.

**Methods:** We used tracking data of bar-headed geese to develop a multi-state movement model and to estimate temporally explicit habitat suitability within the species’ range. We simulated migratory movements between range fragments, and calculated a measure we called route viability. The results are compared to expectations derived from published literature.

**Results:** Simulated migrations matched empirical trajectories in key characteristics such as stopover duration. The viability of the simulated trajectories was similar to that of the empirical trajectories. We found that, overall, the migratory connectivity was higher within the breeding than in wintering areas, corresponding to previous findings for this species.

**Conclusions:** We show how empirical tracking data and environmental information can be fused for meaningful predictions of animal movements throughout the year and even outside the spatial range of the available data. Beyond predicting connectivity, our framework will prove useful for modelling ecological processes facilitated by animal movement, such as seed dispersal or disease ecology.

## Introduction

Understanding the movement of animals is key to understanding the functional connectivity they provide between suitable patches of habitat by transporting biomass, genes, and less mobile organisms. How and when animals move in space, thereby linking areas that are separated in geographical space, has wider ecological implications for the population structure of the species in question, and can also shape the dispersal opportunities for e.g. flowering plants in form of pollen and seeds, or pathogens (Bauer and Hoye, 2014). Establishing connectivity networks and understanding the contribution of animal movement to such networks is a prime motive in ecology, and pivotal to our understanding of spatial structuring processes.

Establishing whether, how, and when animal movement provides a functional connection in space, however, is not easily achieved. Capture-mark-recapture techniques have revealed much about dispersal capabalities of individual animals, thereby providing a history of observed connectivity between distant patches. Estimates such as maximum observed dispersal distances can be used to infer connectivity networks where movement has not been observed, yet there are limitations to their application (Calabrese and Fagan, 2004), as distance alone can be insufficient to explain e.g. patch connectivity. Descriptors of effective rather than e.g. Euclidian distance between patches that incorporate barriers and facilitations to animal movement can be used to improve predictions of connectivity. Algorithms like least-cost paths (e.g., Ferreras, 2001; Graham, 2001) and electrical circuit theory (McRae, Dickson, *et al.*, 2008) can, in combination with spatially explicit predictors of landscape resistance to movement, provide environmentally informed estimates of connectivity between patches (e.g. for population genetics (Row, Blouin-Demers, *et al.*, 2010). Often, however, the animal location data used to inform models used for predicting such resistance surfaces lack a behavioural context, and consequently might not be representative of how animals move through the environment (Keeley, Beier, *et al.*, 2017).

More recently, the equipment of wild animals with remote tracking technology has provided great insights into how, when, and where animals move (Hussey, Kessel, *et al.*, 2015; Kays, Crofoot, *et al.*, 2015). Such data are a rich source of information not only about the movement and behaviour of individuals, but can also reveal the functional, observed connectivity between spatially separated areas in great detail. In combination with environmental information about the utilised habitat, movement data can provide detailed insight into habitat connectivity for the observed individuals (e.g., Almpanidou, Mazaris, *et al.*, 2014). Connectivity estimates derived from observed movement, as e.g. in fragmented landscapes, have been shown to outperform predictions derived from resistance surfaces (e.g., LaPoint, Gallery, *et al.*, 2013). Yet, even though remotely tracked animal movement provides data for the most direct and detailed estimate of connectivity (Calabrese and Fagan, 2004), the use of animal movement data is not without constraints. While the miniaturisation of tracking technologies permits scientists to follow ever more individuals of ever smaller species, the cost and effort associated with animal tracking limit sample size, as well as the spatial and temporal extent of the data that can be collected. Thus, the number of individuals that scientists are realistically able to track will remain minuscule compared to even the most conservative estimates of the numbers of moving animals on this planet. Inferring the movements of unobserved individuals is thus becoming an increasingly important matter to utilise the knowledge from few, well-studied individuals to estimate the behaviour at a population level. However, such generalisations are not straightforward, mainly because the movement behaviour of individuals and the observed variation may not be representative for the population or the whole species (e.g., Austin, Bowen, *et al.*, 2004). Individual decision-making is not only influenced by general species properties, but also variation between individuals and their needs, and the surrounding environmental conditions (Nathan, Getz, *et al.*, 2008). Any kind of movement behaviour is thus to some extent unique to the individual, explicit in time, space, and the environmental conditions as well as specific to the ecological context it happened in.

The literature published on animal movement models is extensive, and such models have been shown to provide useful and sensible estimates on the behaviour of observed as well as unobserved individuals (e.g., Morales, Haydon, *et al.*, 2004; Codling, Plank, *et al.*, 2008; Péron, Fleming, *et al.*, 2017; Michelot, Langrock, *et al.*, 2017). Making such models aware of the environmental context of the movements is however key to providing sensible hypotheses of the routes that animals might take. This requires the contextualisation of observed movement, and the understanding of how animals utilise environmental features for route decision-making. Movement models that incorporate e.g. a resource-selection model (step-selection functions, e.g. Fortin, Beyer, *et al.*, 2005; Thurfjell, Ciuti, *et al.*, 2014) are becoming increasingly popular. Step-selection functions have been shown to yield functional estimates how environmental features influence an animal’s movement through the landscape (e.g., Richard and Armstrong, 2010), and have been used to estimate connectivity between patches (Squires, DeCesare, *et al.*, 2013). Such step-selection functions, representing resource selection during actual movement, can be used to derive behaviour-specific predictions for resistance of a landscape to movement. In combination with least-cost paths or circuit theory, these context-aware resistance surfaces provide the means to predict the movement of individuals through the landscape (e.g., Zeller, McGarigal, *et al.*, 2014; Zeller, McGarigal, *et al.*, 2016).

In many cases, however, animals use series of different movement strategies that change in response to the surrounding environment, or in response to the different needs an animal has for different behaviour or life-history stages. Currently, however, even context-aware approaches used for predicting the movement of unstudied individuals often make the assumption that animals follow a single, constant decision rule. As shown by Zeller, McGarigal, *et al.* (2016), these decision rules can be sensitive to the spatial and temporal scale of the observations, and are considered to be independent of the supply needs of the individual. We think that realistic movement simulations should not only take the environmental context of movement behaviour into account, but also acknowledge the different movement strategies expressed by a species (see e.g. Morales, Haydon, *et al.*, 2004). One example of such a multi-state movement behaviour with striking differences in e.g. spatial scale of movement between states is the stepping-stone-like migrations as performed by many migratory bird species that predominantly use flapping flight for locomotion. Here, we refer to stepping-stone migrations as e.g. performed by large waterbirds like ducks and geese who cover large distances in fast and non-stop flight and use stopover locations for extended staging periods to replenish their fat reserves. Context-aware, multi-state approaches for simulating animal trajectories are however, not easily available. While it is possible to model habitat suitability or resistance or cost surfaces and use modelled movement trajectories to estimate connectivity, the movement and the context represent two separate and static entities. An additional difficulty is that for the formulation of such resistance or cost surfaces detailed a-priori knowledge is required, which again necessitates a level of knowledge that might not be present. In the case of stepping-stone migratory movements, this could refer to a-priori knowledge about available stopover sites for staging migrants.

Here, we would like to introduce a novel approach that allows for inferring environmentally informed migratory trajectories from a multi-state discrete movement model. Using a novel conditional movement model specifically designed for generating random trajectories using template empirical trajectories (Technitis, Weibel, *et al.*, 2016; Technitis, Weibel, *et al.*, in preparation), we developed this approach especially with stepping-stone migrations and similar movement strategies in mind. We extend this movement model to represent the two major states of stepping stone migrations, the non-stop migratory flights and the staging periods, using a stochastic switch informed by empirical estimates of typical duration of both behaviours. Our multi-state movement model can simulate migratory trajectories that are realistic in terms of the geometric properties of empirically collected migratory movements by sampling from empirical distribution functions. We develop a measure of route viability that integrates properties of the simulated trajectory and its environmental context to assess how suitable the simulated migratory route and timing strategy might be for actual, unobserved individuals. For stepping-stone migrations, we here assume that the quality of stopover sites between the breeding grounds and wintering areas predominantly determines how preferable a certain route might be (Green, Alerstam, *et al.*, 2002; Drent, Eichhorn, *et al.*, 2007). While the migration simulation model and the measure of route viability we introduce here are tailored for our study system, we think that this approach in general is flexible and could be applicable to many other study systems and strategies.

Specifically, we apply this approach to a pronounced long-distance migrant: the bar-headed goose (*Anser indicus*, Latham 1790), a Central Asian species of waterbird well known for its incredible performance of crossing the Himalayas during migration. This species has a distribution range that is characterised by four distinct breeding areas, mirrored by four distinct wintering areas south of the Himalayas. The migratory routes for some populations of bar-headed geese are known (e.g., Hawkes, Balachandran, *et al.*, 2011; Guo-Gang, Dong-Ping, *et al.*, 2011; Prosser, Cui, *et al.*, 2011; Bishop, Yanling, *et al.*, 1997; Takekawa, Heath, *et al.*, 2009): Previous tracking studies have revealed that large parts of the respective populations migrate from their breeding grounds in Mongolia, northern China and on the Tibetan Plateau over the Himalayas to their wintering grounds on the Indian subcontinent. But while the crossing of the Himalayas has been studied in great detail (Hawkes, Balachandran, *et al.*, 2011; Hawkes, Balachandran, *et al.*, 2013; Bishop, Spivey, *et al.*, 2015), less is known about the connectivity between range fragments both within the wintering and within the breeding range (Takekawa, Heath, *et al.*, 2009). The bar-headed goose thus provides a suitable study species for our approach. We will establish a model for bar-headed goose migrations from previously published tracking data, and simulate migrations of unobserved individuals between all fragments of the species’ distribution range. We will assess the viability of these trajectories during several times of year using a segmented habitat suitability model to derive a dynamic migratory connectivity network. To test whether this migratory connectivity network could serve as a quantitative null hypothesis for bar-headed goose migration, we will test our predictions against two very simple hypotheses generated from previously published studies.

Stable isotope analyses suggested that the connectivity within the breeding range of bar-headed geese is relatively high (Bridge, Kelly, *et al.*, 2015), a notion that has been supported by tracking data as well (Cui, Hou, *et al.*, 2010). In the wintering range, however, relatively few movements have been observed (Kalra, Kumar, *et al.*, 2011). Based on these findings (Bridge, Kelly, *et al.*, 2015; Kalra, Kumar, *et al.*, 2011; Cui, Hou, *et al.*, 2010), we expect to find a higher overall viability of trajectories between the fragments of the breeding range than within the wintering range. We further predict that on average, the temporal variation in plausibility of simulated migratory routes within the breeding grounds should be higher than within the wintering grounds. Overall, we would like to introduce a new approach for deriving environmentally informed quantitative null hypotheses for animal movement which can be utilised for estimating migratory connectivity based on limited observations (summarised in Figure 1).

**Figure 1.**
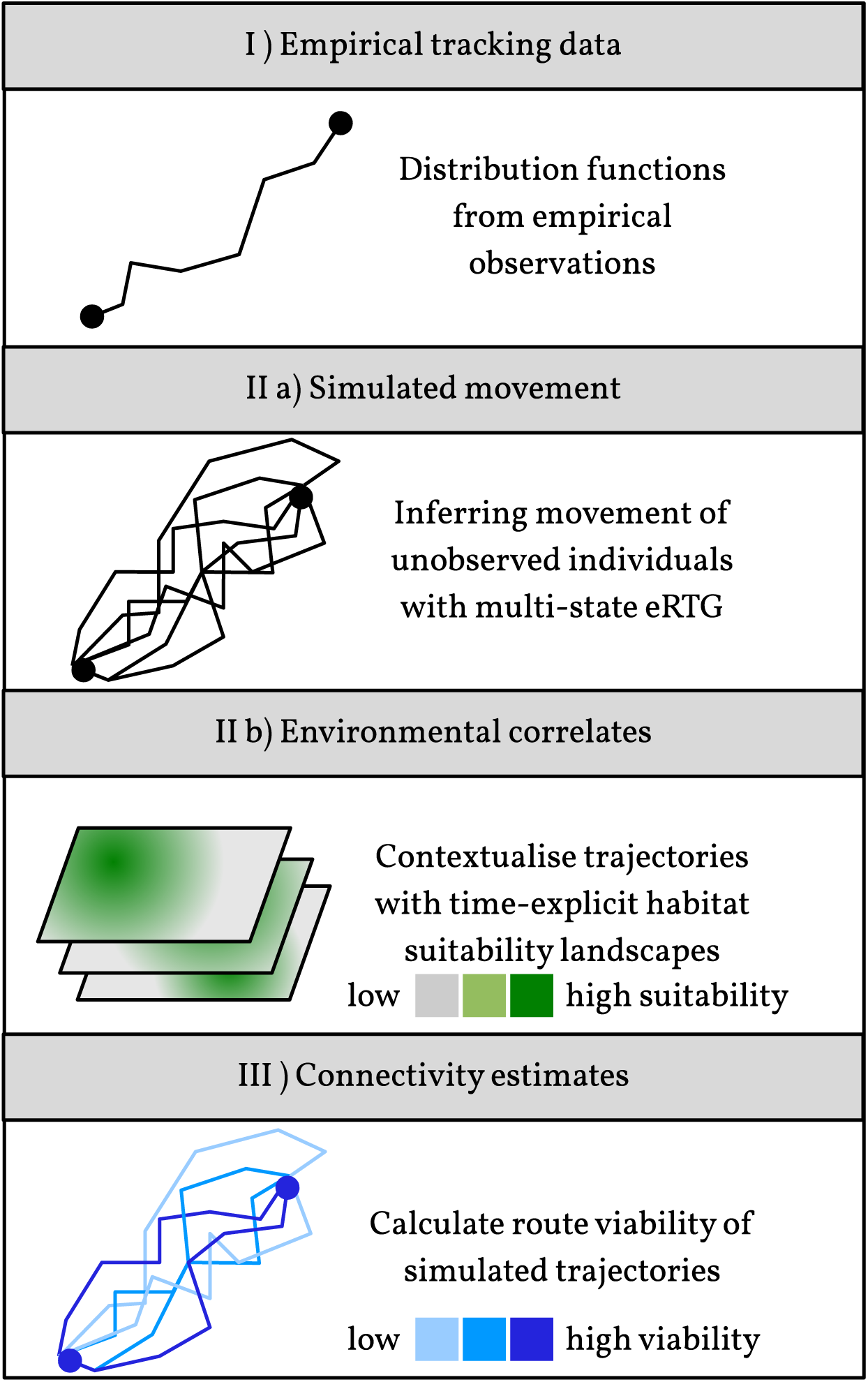
General concept for our approach of environmentally informing simulated stepping-stone migrations: I) Empirical tracking data are IIa) used to derive an informed eRTG to simulate conditional movement between sites of interest, and IIb) combined with environmental correlates to derive predictions of relevant measurements of landscape permeability (here: suitability of stopover sites). III) Finally, the simulated conditional trajectories are evaluated based on characteristics of the trajectory and permeability using an informed measure of route viability.

## Methods

### Tracking data and movement model

Tracking data of bar-headed geese were available to us from a broader disease and migration ecology study implemented by the Food & Agriculture Organization of the United Nations (FAO) and United States Geological Survey (USGS). In total, 91 individuals were captured during the years 2007-2009 in several locations: Lake Qinghai in China (hereafter termed “Lake Qinghai”), Chilika Lake and Koonthankulum bird sanctuary in India (hereafter termed “India”), and Terkhiin Tsagaan Lake, Mongolia (hereafter termed “West Mongolia”). All individuals were equipped with ARGOS-GPS tags which were programmed to record the animals’ location every two hours (ARGOS PTT-100; Microwave Telemetry, Columbia, Maryland, USA). The tags collected and transmitted data for 241 ± 253 (mean ± s.d.) days, and in total 169′887 fixes could be acquired over the course of the tracking period (see also Table 1 & Hawkes, Balachandran, *et al.*, 2011; Takekawa, Heath, *et al.*, 2009). Individuals that were tracked for less than a complete year were excluded from the subsequent analyses, which left a total of 66 individuals (Lake Qinghai: 20, India: 20, West Mongolia: 26). We pooled data from all capture sites for the analyses.

**Table 1.** A summary of the catching sites and corresponding sample sizes. The number of tracking days and GPS fixes are listed as a mean per individual.

| **Capture site** | **Year of capture** | **Sample size**<br>(individuals) | **First fix taken** | **Tracking days**<br>median;<br>[25%; 75% quantile] | **GPS fixes**<br>median<br>[25%; 75% quantile] |
| --- | --- | --- | --- | --- | --- |
| Lake Qinghai | 2007 | 13 | Mar 25 – 31 | 303 [207; 411] | 1670 [682; 2565] |
|  | 2008 | 10 | Mar 30 – Apr 4 | 396 [260; 845] | 2211 [1341; 3573] |
| India | 2008 | 17 | Dec 10 – 18 | 129 [92; 401] | 2060 [1578; 2714] |
|  | 2009 | 7 | Jan 27 – Feb 06 | 134 [53; 448] | 1321 [1107; 3800] |
| West Mongolia | 2008 | 19 | Jul 13 – 15 | 122 [90; 190] | 537 [366; 1312] |
|  | 2009 | 14 | Jul 05 – 08 | 105 [100; 128] | 421 [330; 473] |

We used the recently developed the **e**mpirical **R**andom **T**rajectory **G**enerator (**eRTG**, Technitis, Weibel, *et al.*, 2016; Technitis, Weibel, *et al.*, in preparation) to simulate the migrations of unobserved individuals of bar-headed geese. This movement model is conditional, i.e. simulates the movement between two end locations with a fixed number of steps based on a dynamic drift derived from a step-wise joint probability surface. One main advantage of the eRTG is that the trajectories it simulates retain the geometric characteristics of the empirical tracking data (step length, turning angle, as well as covariance and auto-correlation of step length and turning angle), as it relies entirely on empirical distribution functions. Consequently, if a destination cannot be reached within the realms of the empirical distributions of e.g. step lengths and turning angles, the simulation fails rather than forcing the last step towards the destination.

We extended this movement model by incorporating a stochastic switch between the two main states of bar-headed goose migration, non-stop migratory flights (“*migratory state*”) and movements during staging periods at stopover locations (“*stopover state*”). We classified the entire tracking data according to the individuals’ movement behaviour to identify these states prior to extracting the empirical distributions functions for the eRTG. First, we clustered the locations in the tracking data into four behavioural classes (slow speed & low turning angles, slow speed & high tuning angles, high speed & low turning angles, and high speed & high turning angles) using an expectation-maximisation binary clustering algorithm designed for annotating animal movement data (EMbC, Garriga, Palmer, *et al.*, 2016). We then re-classified the tracking data into two behavioural classes, namely high-speed movements (combining the two high speed classes) and low-speed movements (combining the two low speed classes). Within the high-speed behavioural cluster, the average speed between locations was 8.4±6.7*m*/*s* (mean ± s.d.) whereas the average speed for the low-speed behavioural cluster was 0.3±1.0*m*/*s* (mean ± s.d.). As estimates of speed and turning angle are highly dependent on the sampling rate of the data, we removed those parts of the trajectories that exceeded the average sampling interval of two hours. Subsequently, we used the low-speed locations for the empirical distribution functions of the staging period of the two-mode eRTG, and the locations classified as high-speed for the empirical distribution functions for the migratory component of the eRTG (see Figure S2). Finally, we derived the step lengths and turning angles from each coherent stretch of data (i.e. only subsequent fixes with a sampling rate of 2 hours). Following this, we calculated the changes in step length and turning angle at a lag of one observation, as well as the covariance between contemporary observations of step length and turning angle. We derived the corresponding empirical distribution functions for both movement states and prepared them for use in the eRTG functions.

Finally, we determined the duration of staging periods, and the duration and cumulative distance of individual migratory legs from the tracking data. We first identified seasonal migration events between breeding and wintering grounds (and vice versa) in the empirical trajectories using the behavioural annotation. We then determined migratory legs (sequential locations classified as migratory state) as well as stopovers (sequential locations classified as stopover state, with a duration > 12*h*). We used two main proxies to characterise migratory legs, namely cumulative migratory distance as well as duration, and one proxy to characterise staging periods, namely stopover duration. We calculated these proxies for all individuals and migrations, and determined the maximum observed distance (*dm*_max_) and duration (*Tm*_max_) of a migratory leg. As we did not distinguish between extended staging (e.g. during moult, or after unsuccessful breeding attempts) from use of stopover locations during migration, we calculated the 95% quantile of the observed stopover durations (*Ts*_max_) rather than the maximum.

#### Simulating a bar-headed goose migration with the two-state eRTG

When simulating a conditional random trajectory between two arbitrary locations *a* and *z*, the two-state eRTG initially draws from the distribution functions for the migratory state, producing a fast, directed trajectory. To determine the time available for moving from *a* to *z*, we assumed the mean empirical flight speed derived for the migratory state, and calculated the number of required steps accordingly. While simulating the trajectory, after each step modelled by the eRTG, the cumulative distance of the trajectory (*dm*) as well as the duration (*Tm*) since the start of the migratory leg were calculated. By using *dm*, *Tm*, as well as the empirically derived *dm*_max_ and *Tm*_max_, our two-state eRTG was based on a binomial experiment with two possible outcomes: switching to the stopover state with a probability of *p*_*ms*_, or resuming migration with a probability of 1 – *p*_*ms*_. We defined *p*_*ms*_, the transition probability to switch from migratory state to stopover state, as

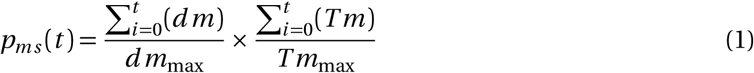

At step *t*, the simulation of the migratory movement can switch to the unconditional stopover state, corresponding to a correlated random walk, with a probability of *p*_*ms*_(*t*). Likewise, the simulation can switch back from stopover state to migratory state with the probability *p*_*sm*_(*t*), which we defined as as

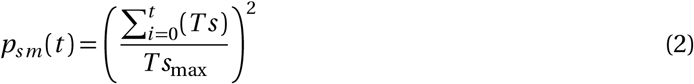

This process is then repeated until the simulation terminates because: either the trajectory reached its destination, or the step-wise joint probability surface did not allow for reaching the destination with the remaining number of steps (resulting in a dead end or zero probability).

### Evaluating the plausibility of simulated migrations

We estimated the plausibility of each simulated trajectory representing a unique migratory route, using a measure we called route viability Φ aimed to integrate the ecological context into the movement simulations. We developed this measure specifically with the stepping stone migratory strategy of bar-headed geese or similar species in mind, and it is defined by the time spent in migratory mode, the time spent at stopover sites, and the habitat suitability of the respective utilised stopover sites. For this specific measure of route viability, we make two main assumptions: (1), it is desirable to reach the destinations quickly, i.e. staging at a stopover site comes at the cost of delaying migration, and (2), the cost imposed by delaying migration is inversely-proportional to the quality of the stopover site, i.e. the use of superior stopover sites can counterbalance the delay. Our argumentation for these assumptions is that during spring migration, the arrival at the breeding grounds needs to be well-timed with the phenology of their major food resources (Bauer, Gienapp, *et al.*, 2008). Furthermore, the quality of stopover sites has been shown to be of crucial importance for other species of geese with similar migratory strategies (Green, Alerstam, *et al.*, 2002; Drent, Eichhorn, *et al.*, 2007).

Each simulated multi-state trajectory between two arbitrary locations *a* and *z* can be characterised by a total migration duration *τ*_*a, z*_, which consists of the total flight time *τ*_*M, a, z*_ and the total staging time at stopover sites *τ*_*S, a, z*_. The total flight time *τ*_*M, a, z*_ is the sum of the time spent flying during each migratory leg *l*, and is thus 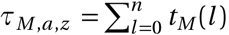 with *t*_*M*_(*l*) corresponding to the time spent flying during migratory leg *l*. Similarly, the total staging time *τ*_*S, a, z*_ consists of the staging times at all visited stopover sites, corresponding to 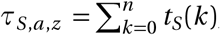, where *t*_*S*_(*k*) amounts to the staging time at stopover site *k*. For our metric of route viability, we will consider the time spent staging at stopover locations *τ*_*S, a, z*_ as a delay compared to the time spent in flight. This delay is, however, mediated by the benefit *b* an individual gains at the stopover site, e.g. by replenishing its fat reserves. We define that this benefit gained by staying at a stopover site *k*, *b*(*k*), is proportional to the time spent at site *k*, *t*_*S*_(*k*), and the habitat suitability of site *k*, *S*(*k*). This habitat suitability *S* should range between [0, 1], which allows our measure of route viability to range between [0, 1] as well. We further assume the effects of several sequential stopovers to be cumulative, and thus define the total benefit of a migratory trajectory between locations *a* and *z* with *n* stopovers as 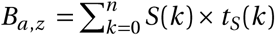. Finally, we define the route viability Φ_*a, z*_ of any trajectory between *a* and *z* as:

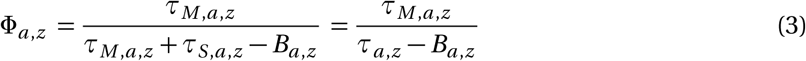

In this way, the viability of a trajectory with no stopovers and a trajectory with stopovers of the highest possible quality (*S*(*k*) = 1) will be equal, and is defined solely by the time the individual spent in migratory mode (Φ_*a, z*_ = 1). For trajectories with stopovers in less than optimal sites, however, the viability of trajectories is relative to both the staging duration and quality of stopover sites, and should take values of 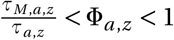. Using this metric, we assessed simulated trajectories in a way that is biologically meaningful for bar-headed geese. In the next section we detail how we calculated the route viability Φ for each simulated migration.

### A network of migratory connectivity for bar-headed geese migrations

We simulated migrations of bar-headed geese within the native range of the species which naturally occurs in Central Asia (68–107^°^*N*, 9–52^°^*E*). According to BirdLife International and NatureServe (2013), both the breeding and wintering range are separated into four distinct range fragments (see also Figure S1), with minimum distances between range fragments ranging from 79*km* to 2884*km*. For this study, we investigated how well, in terms of an environmentally informed measure of route viability and e.g. the number of stopovers required to reach a range fragment, these range fragments can be connected by simulated migrations of bar-headed geese.

To choose start-and endpoints for the simulated migrations, we sampled ten random locations from each of these range fragments indicated in the distribution data provided by BirdLife International and NatureServe (2013). We simulated 1000 trajectories for all pairs of range fragments (100 trajectories per location pair) and counted the number of successes (trajectories reach the destination) and failures (trajectories terminate in a dead end). We then assessed all successful trajectories by counting how many stopover locations were utilised in each trajectory. We proceeded to calculating the viability of simulated routes in the following way: Initially, we determined the total duration of the migration of trajectory between locations *a* and *z*, *τ*_*a, z*_, the number of stopover sites used, *n*_*a, z*_, as well as the time spent at each stopover site (*t*_*S*_(*k*)) for each of a total of the *n*_*a, z*_ stopovers (corresponding to the number of steps multiplied with the location interval of two hours). We determined the habitat suitability of stopover locations *S*(*k*) using habitat suitability landscapes that represented habitat suitability for bar-headed geese during five periods of the year (see Figure S3): winter/early spring (mid-November - February), mid-spring (mid March - mid April), late spring/summer (mid April - mid August), early autumn (mid August-mid September), and late autumn (mid September - mid November). We identified these periods using a segmentation by habitat use (van Toor, Newman, *et al.*, 2016, for details see Section A in the Electronic Supplementary Material (ESM)). The segmentation by habitat use uses animal location data and associated environmental information to identify time periods for which habitat use is consistent. Habitat suitability models derived for these time periods should thus reflect differences in habitat use by bar-headed geese throughout the year. We used time series of remotely sensed environmental information and random Forest models (Breiman, 2001) to derive habitat suitability models corresponding to these five time periods, and predicted the corresponding habitat suitability landscapes (section A in the ESM). Following the prediction of habitat suitability landscapes for winter/early spring, mid-spring, late spring/summer, early autumn, and late autumn, we annotated the all stopover state locations of the simulated trajectories with the corresponding habitat suitability. We then calculated the benefit *b* gained by using a stopover location *k* using the mean suitability for each of the stopover locations, *S*(*k*), and the duration spent at stopover locations, *τ*_*S*_(*k*).

To calculate the route viability Φ_*a, z*_, we also required an estimate for duration of migration if a simulation were exclusively using the migratory state *τ*_*M, a, z*_, without the utilisation of stopover sites. We used a simple linear model to predict flight time as a function of geographic distance which we trained on the empirical data derived from the migratory legs (see Section B in the ESM for details). By basing the linear model on the empirical migratory legs rather than mean flight speed, the estimate for *τ*_*M*_ retains the inherent tortuosity of waterbird migrations. For each simulated trajectory, we then calculated the geographic distance between its start-and endpoint, and predicted the expected flight time *τ*_*M, a, z*_. Finally, we calculated route viability Φ_*a, z*_ for all trajectories using equation 3, repeating the process for every of the five suitability landscapes derived from the segmentation by habitat use. This resulted in five different values of Φ_*a, z*_ for every simulated trajectory, corresponding to winter/early spring, mid-spring, late spring/summer, early autumn, and late autumn, respectively.

#### Calculating migratory connectivity as average route viability

We calculated migratory connectivity between range fragments as the average route viability Φ_avg._ of all trajectories connecting two range fragments. We calculated this average by using non-parametric bootstrapping on the median route viability Φ_avg._ (using 1000 replicates), and also computed the corresponding 95% confidence intervals (CI) of the median route viability Φ_avg._. We did this for each of the five time periods represented in the suitability landscapes, and also computed an overall migratory connectivity by averaging all five habitat suitability values for each stopover site prior to calculating Φ.

We wanted to compare migratory connectivity within the breeding range and migratory connectivity in the wintering range to test our first hypothesis stating that migratory connectivity should be higher within the breeding range. To do so, we differentiated between route viability among breeding range fragments (Φ_*breeding*_), among the wintering range (Φ_*wintering*_), and between breeding and wintering range fragments (Φ_*mixed*_). We computed the median and 95% CIs of route viability with non-parametric bootstrapping with 1000 replicates, using the average habitat suitability of all five suitability landscapes for all trajectories within the breeding range, all trajectories in the wintering range, and all trajectories connecting breeding range fragments with wintering range fragments.

To test our second hypothesis, stating that variation in migratory connectivity throughout the year should be higher in the breeding range than in the wintering range, we calculated the standard deviation of route viability for the five suitability landscapes in the breeding range and in the wintering range. We did this by again, by differentiating trajectories in the wintering range, trajectories in the breeding range, and trajectories connecting breeding range fragments with wintering range fragments. We computed route viability Φ for each of the five suitability landscapes for all trajectories, and pooled the corresponding values for Φ_late winter/early spring_, Φ_mid-spring_, Φ_late spring/summer_, Φ_early autumn_, and Φ_late autumn_ for the wintering range, for the breeding range, and for trajectories connecting breeding range fragments with wintering range fragments separately. We then used a non-parametric bootstrapping (1000 replicates) on the standard deviation over the five time periods, and determined the corresponding 95% CIs on the standard deviation.

#### Calculating route viability for empirical migrations

Following this, we annotated the stopover locations of empirical migrations with the habitat suitability of the corresponding time period, and calculated the route viability for these migratory trajectories in the same way as described above. We then used non-parametric bootstrapping on the median route viability for all empirical migrations (Φ_emp., total_), only spring migrations (Φ_emp., spring_) and only autumn migrations (Φ_emp., autumn_), and computed 95% CIs for the median of Φ_emp., total_, Φ_emp., spring_, and Φ_emp., autumn_.

## Results

### Route viability of empirical and simulated migrations

The simulations resulted in a total of 30730 simulated trajectories, of which 8945 trajectories connected breeding range fragments (simulation success rate: 74.5%), 5393 trajectories connected wintering range fragments (simulation success rate: 44.9%), and the remaining 16392 trajectories connected breeding and wintering range fragments (simulation success rate: 51.2%; see Figure S4). While all these trajectories were successful in connecting origin and destination (i.e. did not result in a dead end), they differed profoundly in their route viability Φ_simulated_, which ranged between 0.014 and 0.59. We found that simulated migrations had a higher route viability for late spring and summer than for autumn (Figure 2).

**Figure 2.**
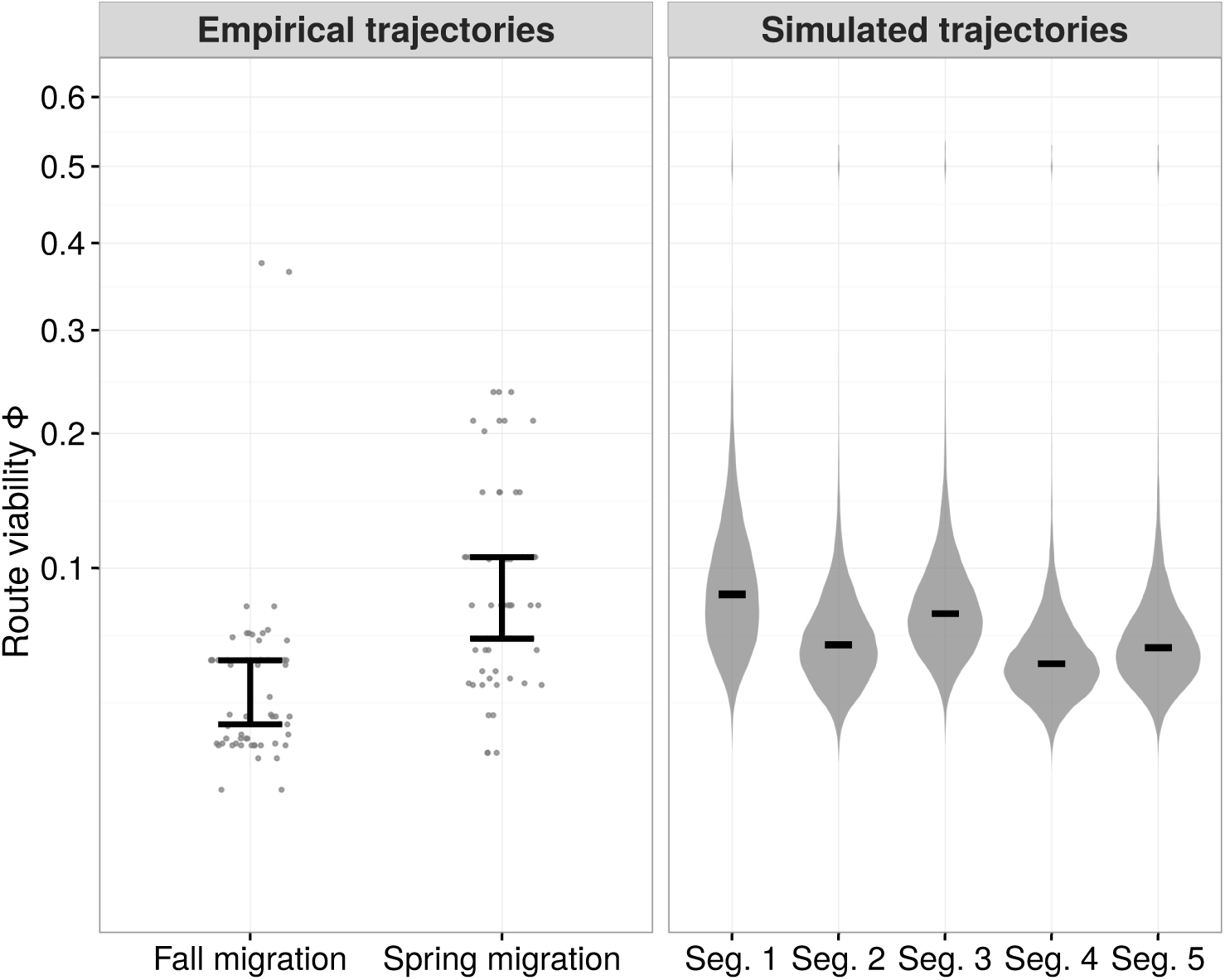
The route viability Φ of empirical and simulated migrations. Here we show Φ for spring and autumn migrations, as well as the Φ for the simulated trajectories across all five suitability landscapes (Seg. 1: winter/early spring, Seg. 2: mid-spring, Seg. 3: late spring/summer, Seg. 4: early autumn, Seg. 5: late autumn). The black bars show the 95% CIs for the respective medians, and the grey dots and violin plots show the observed (empirical trajectories) and densities for the observed route viability (simulated trajectories).

The range of route viability for simulated migrations was comparable to that of the empirical migrations (Φ_emp., total_: 0.01 – 0.38). Overall, we found that route viability of empirical migrations was higher for spring migrations (Φ_emp., spring_: [0.0614;0.1070]; 95% CIs on the median) than for autumn migrations (Φ_emp., autumn_: [0.0270;0.0514]; 95% CIs on the median). This was caused both by differences in the habitat suitability of utilised stopover locations and the differences in migration duration between spring and autumn migrations. We found that bar-headed geese on average stayed longer at stopover locations during autumn than during spring migrations (spring: 6.8±14.2 days, autumn: 11.8±12.2 days; mean ± s.d.).

### Route viability of simulated migrations in the species’ range

We separated the simulated trajectories into movements within the breeding range, movements within the wintering range, and movements resembling seasonal migrations between the breeding and wintering range. Here, we found that viability of trajectories was highest within the breeding range (95% CIs for median Φ_breeding_: [0.0676;0.0684]; 95%-quantiles of median Φ_breeding_: [0.1469;0.1546]), and lowest within the wintering range (95% CIs for median Φ_wintering_: [0.0590;0.0596]; 95%-quantile of median Φ_wintering_: [0.1090;0.1147]), predicting that movements between range fragments should occur more often within the breeding than in the wintering areas. The median route viability for migrations between breeding and wintering range fragments was intermediate (95% CIs for median Φ_mixed_: [0.0618;0.0622]; 95%-quantile of median Φ_mixed_: [0.1224;0.1296]). These patterns are reflected in the simplified network of average migratory connectivity Φ_avg._ (Figure 3). We also identified the single trajectory with the maximum route viability between range fragments rather than the median (Figure S5). This network of maximum migratory connectivity shows that migrations that connect the breeding and wintering ranges have the highest route viability. Finally, the number of stopover locations of movements was proportional to the geographic distance between range fragments (Figure S6).

**Figure 3.**
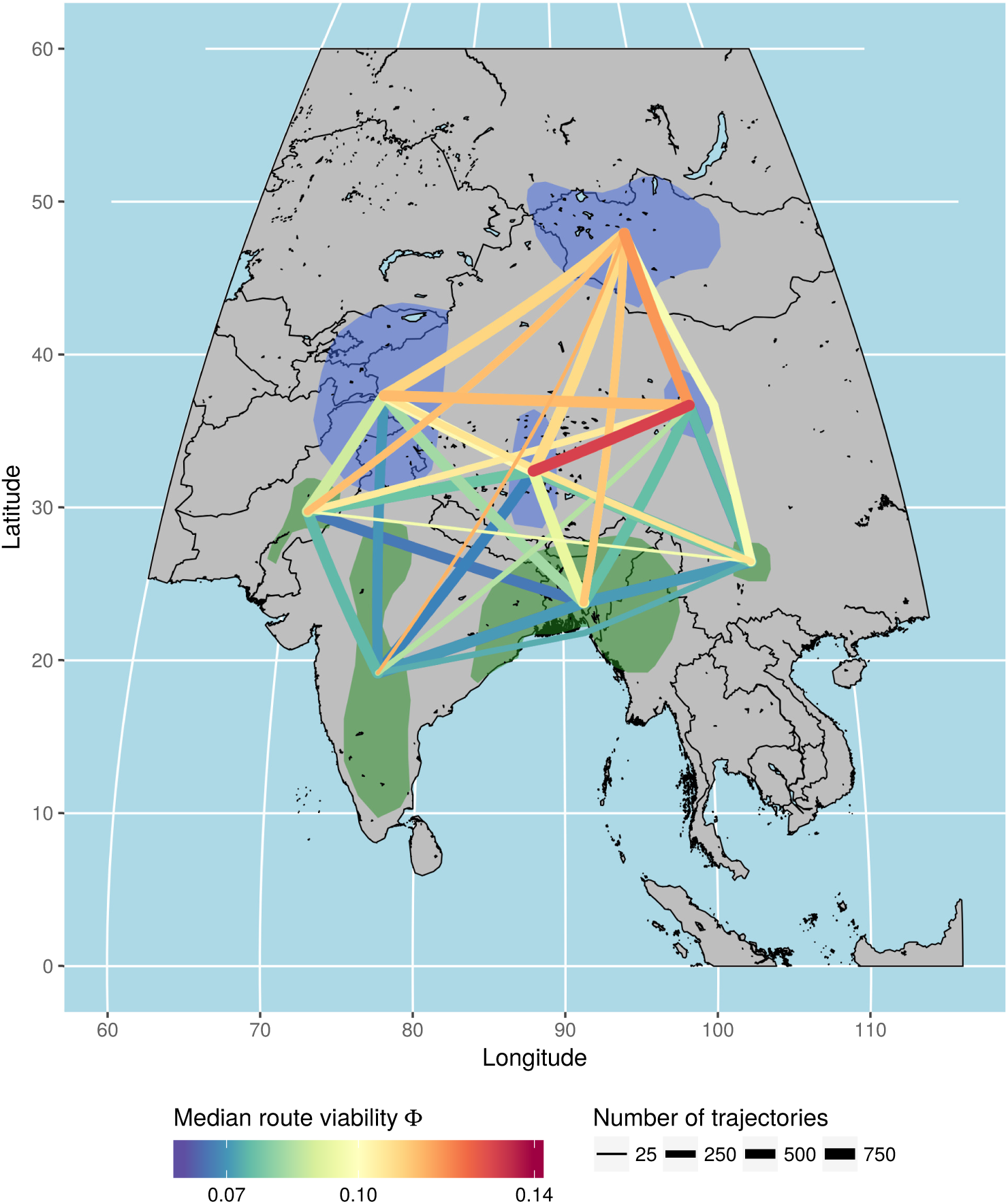
The median route viability Φ between range fragments of bar-headed geese. We summarised Φ for all pairwise range fragment trajectories using the median route viability. The thick-nessofedges represents the sample size. Bluepolygonsshow thenative breeding areaofthe species, green polygons the native wintering range. Long edges are curved for sake of visibility.

### Temporal variability of within-range migratory connectivity

We found that the spatial patterns of migratory connectivity varied across the suitability landscapes derived from the five habitat suitability landscapes representing five periods of consistent habitat suitability (Figure 4; see also Figure S3 for details on the temporal correspondence of the time periods). For the suitability landscapes derived for winter/early spring, mid spring, and late spring/summer, the estimated connectivity predicts that bar-headed goose migrations are most likely to occur between the wintering and breeding range, and within the breeding range. For early autumn, connectivity patterns predict that movement should be most likely between breeding and wintering areas. For late autumn, we also observed that connectivity predicts movement within the wintering range of the species. We also calculated the 95% CIs for the overall migratory connectivity values for each time period (Figure 2), which predicts the highest median route viability for the periods from winter/early spring (mid-November - February) as well as from late spring/summer (mid April - mid August). We also compared the standard deviation of route viability across suitability landscapes and found the highest variation for the breeding range (95% CIs for s.d. of Φ_breeding_: [0.0124;0.0133]) and the lowest variation for the wintering range (95% CIs for s.d. of Φ_wintering_: [0.0041;0.0046]). Again, the trajectories between breeding and wintering range fragments showed intermediate values (95% CIs for s.d. of Φ_mixed_: [0.0084;0.0089]).

**Figure 4.**
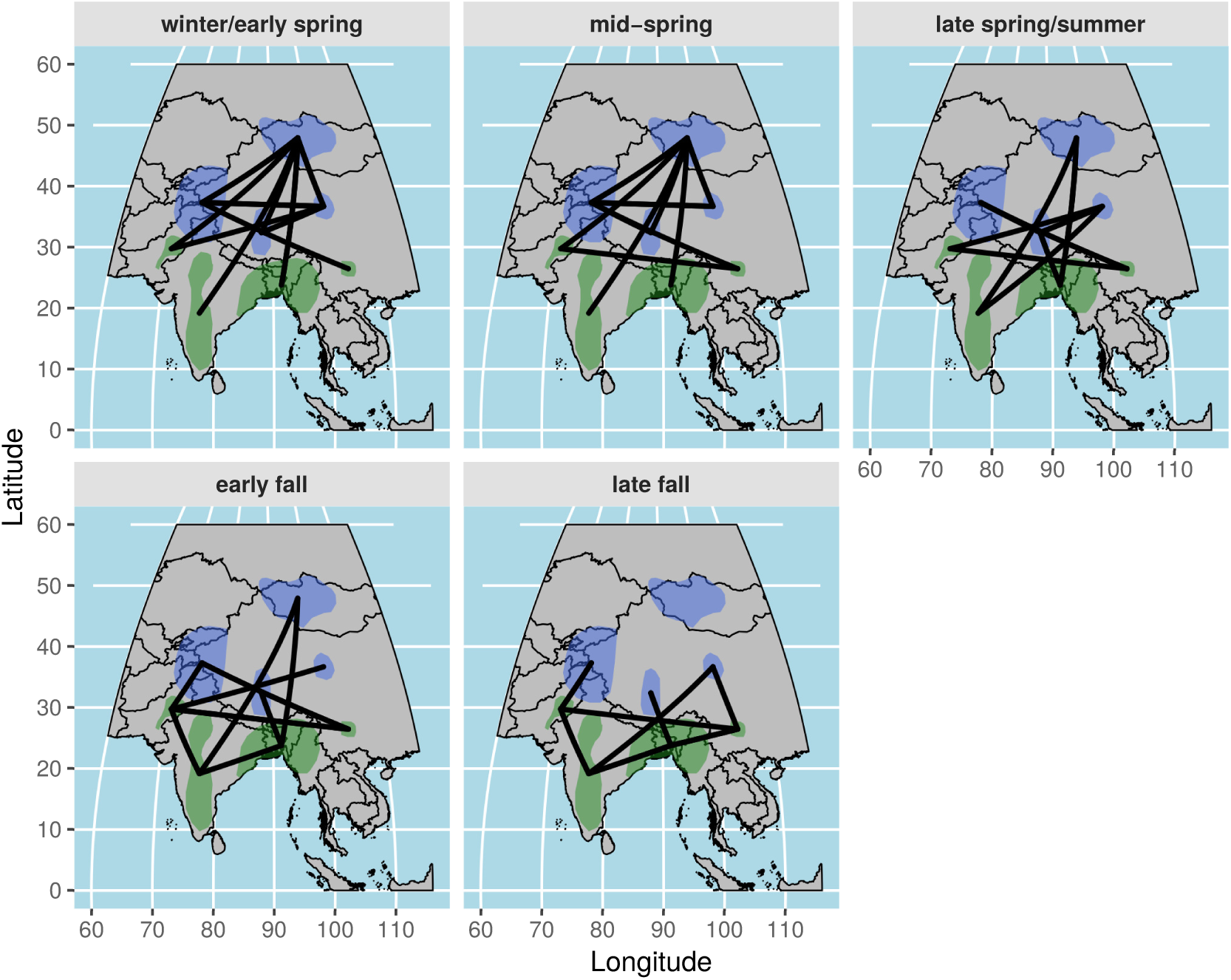
This Figure Shows the temporal dynamics of the route viability Φ by showcasing the predicted movements for each of the five suitability landscapes separately. The visible edges of the network have a median route viability Φ that is higher than 75% of the ecological likelihood for the complete network. The respective time periods associated to these networks is displayed in Figure S3.

## Discussion & Conclusions

Using previously collected tracking data of bar-headed geese and the novel empirical Random Trajectory Generator, we were able to successfully develop a model that can simulate the migratory movements of bar-headed geese. Our extension of the eRTG with a stochastic switch between a migratory state and a stopover state was sufficient to capture the overall migratory strategy of this species. With this model for bar-headed goose migrations, we inferred the migrations of unobserved individuals between all fragments of the species’ distribution range, and used an environmentally informed measure of route viability to derive average estimates of migratory connectivity between range fragments. We put this simplified predictive network of migratory connectivity to a simple test using predictions derived from the literature. Indeed, we found that the average route viability, as an indicator of migratory connectivity, was higher within the species’ breeding range (Φ_breeding_) than in the wintering areas Φ_wintering_, confirming the expectations from the literature (Cui, Hou, *et al.*, 2010; Kalra, Kumar, *et al.*, 2011; Bridge, Kelly, *et al.*, 2015). While bar-headed geese are thought to be philopatric to their breeding grounds (Takekawa, Heath, *et al.*, 2009), the post-breeding period seems to be a time of great individual variability and extensive movements (Cui, Hou, *et al.*, 2010). This has also been observed for other Anatidae species (e.g., Gehrold, Bauer, *et al.*, 2014), as due to the temporary flightlessness during moult the choice of suitable moulting sites is critical to many waterfowl species. As the average route viability within the breeding range and during the summer months is high, we think that unsuccessfully breeding bar-headed geese and individuals in the post-breeding period are not limited by sufficiently suitable stopover locations when moving between breeding range fragments. Furthermore, our results confirmed that the temporal variability of migratory connectivity was higher in the breeding areas north of the Himalayas than in the subtropical wintering areas.

We think that simulating trajectories with multiple movement states and an element of randomness can be useful to infer the movements of unobserved individuals. Here, we simulated migratory trajectories under the assumption that the movements of the tracked individuals are similar to those of other individuals. While this limits the informative value for an estimate of migratory connectivity, we think that through repeated simulations, it is possible to explore the routes of unobserved individuals according to the movement behaviour observed in empirical data on the same temporal scale. Indeed, average route viability corroborated previous studies on the within-range movements for bar-headed geese even without additional filtering. Consequently, we think that in combination with relevant environmental and ecological information, the simulation of unobserved migrations using a model like our two-state eRTG can provide a sensible and quantitative null hypothesis for the migrations of bar-headed geese or species with similar strategies. While we determined the route viability using only the habitat suitability of the stopover locations, apart from the some measures of migratory duration, we think that other correlates such as wind support or altitude profile could easily be incorporated for the migratory state. Similarly, the transition probabilities that mediate the switch between movement modes can be extended to include environmental conditions. In general, we think that our stochastic switch performed reasonably well in replicating the movement behaviour observed from recorded tracks. We used simple functions to determine transition probabilities due to the gappy nature of the data and long fix interval (two hours). If a larger sample size were available, the functions we used (see equations 1 and 2) could be replaced by a probability distribution function that more adequately represents the decision-making of bar-headed geese. Alternatively, algorithms such as state-space models could be integrated to simulate animal movement with a more complex configuration of movement states (Morales, Haydon, *et al.*, 2004; Patterson, Thomas, *et al.*, 2008). Overall, we think that with modifications specific to the species of interest, the approach described in this study could be adapted for other scenarios of animal movement. One important application for our or similar approaches could be to support capture-mark-recapture data, especially when tracking data for multiple individuals are hard to acquire. Simulations from a multi-state movement model informed by the movements of few representative individuals could be used to infer alternative routes connecting the re-sightings of individually marked animals. A corresponding relevant measure of route viability could then be used to explore the alternative strategies from an ecologically informed perspective. In such a study, it could also be of interest to use Bayesian approaches to approximate ideal routes using the environmental context.

Furthermore, we think that our results highlight the importance of integrating temporal changes in habitat use of moving animals into measures of landscape connectivity. This was already pointed out by Zeigler and Fagan (2014), who argued that the ecological function of landscape connectivity through animal movement is not only determined by *where*, but also *when* the environment provides the conditions that allow an individual to move from *a* to *z*. In our study, estimates of migratory connectivity were affected by changes in the predicted habitat suitability of stopover locations, whereas in other cases, changes in vegetation density throughout the year or the temporary freezing of waterbodies can be imagined to change connectivity between distant sites. Using time series of environmental information in combination with an approach segmenting a species utilisation of the environment for moving, as shown here, could help with the identification of temporal patterns of landscape connectivity. Accounting for such temporal changes in connectivity could also help better understand how for example diseases can spread through through a metapopulation (e.g. white-nose syndrome, Blehert, Hicks, *et al.*, 2009; Turner, Reeder, *et al.*, 2011, or Influenza A viruses in birds, Gaidet, Cappelle, *et al.*, 2010; Newman, Hill, *et al.*, 2012).

Overall, models that incorporate a species’ movement behaviour and its utilisation of the environment can provide sensible estimates for landscape connectivity, and possibly for a wider range of applications. We think that our approach provides a starting point for complementing tracking efforts with ecologically relevant estimates of a species’ potential to migrate through a landscape and act as a link between patches, populations, and ecosystems.

## Acknowledgements

This work was made possible by the efforts of the many cooperating scientists in China, India, and Mongolia who assisted the Wildlife Health and Ecology Program at the Food and Agriculture Organization of the United Nations (FAO) and US Geological Survey (USGS) in the collection of the tracking data with ecological findings provided in other reports and papers. We are indebted to Markus Rampp and Karin Gross at the Computation Center of the Max Planck Society for their support with the computational facilities. The use of trade, product, or firm names in this publication is for descriptive purposes only and does not imply endorsement by the US Government or FAO. The views expressed in this publication are those of the author(s) and do not necessarily reflect the views or policies of the FAO or the US Goverment. This study re-analysed several datasets from FAO and USGS published studies; Institutional Animal Care and Use Committee details are available in the original publications. MLvT was supported by the International Max Planck Research School for Organismal Biology and the European Union’s Horizon 2020 research and innovation programme under grant agreement No 727922 (Delta-Flu).

## References

Almpanidou, V, Mazaris, AD, Mertzanis, Y, et al. (2014). Providing insights on habitat connectivity for male brown bears: A combination of habitat suitability and landscape graph-based models. Ecological Modelling 286:37–44.

Austin, D, Bowen, WD, and McMillan, JI (2004). Intraspecific Variation in Movement Patterns: Modeling Individual Behaviour in a Large Marine Predator. Oikos 105:15–30.

Bauer, S and Hoye, BJ (2014). Migratory Animals Couple Biodiversity and Ecosystem Functioning Worldwide. Science 344:1242552.

Bauer, S, Gienapp, P, and Madsen, J (2008). The Relevance of Environmental Conditions for Departure Decision Changes En Route in Migrating Geese. Ecology 89:1953–1960.

BirdLife International and NatureServe (2013). Bird Species Distribution Maps of the World. Version 3.0.

Bishop, CM, Spivey, RJ, Hawkes, LA, et al. (2015). The Roller Coaster Flight Strategy of Bar-Headed Geese Conserves Energy during Himalayan Migrations. Science 347:250–254.

Bishop, MA, Yanling, S, Zhouma, C, et al. (1997). Bar-Headed Geese *Anser Indicus* Wintering in South-Central Tibet. Wildfowl 48:118–126.

Blehert, DS, Hicks, AC, Behr, M, et al. (2009). Bat White-Nose Syndrome: An Emerging Fungal Pathogen? Science 323:227–227.

Breiman, L (2001). Random Forests. Mach Learn 45:5–32.

Bridge, ES, Kelly, JF, Xiao, X, et al. (2015). Stable Isotopes Suggest Low Site Fidelity in Bar-Headed Geese (*Anser Indicus*) in Mongolia: Implications for Disease Transmission. Waterbirds 38:123–132.

Calabrese, JM and Fagan, WF (2004). A comparison-shopper’s guide to connectivity metrics. Frontiers in Ecology and the Environment 2:529–536.

Codling, EA, Plank, MJ, and Benhamou, S (2008). Random Walk Models in Biology. J R Soc Int 5:813–834.

Cui, P, Hou, Y, Tang, M, et al. (2010). Movement Patterns of Bar-Headed Geese *Anser Indicus* during Breeding and Post-Breeding Periods at Qinghai Lake, China. J Ornithol 152:83–92.

Drent, RH, Eichhorn, G, Flagstad, A, et al. (2007). Migratory Connectivity in Arctic Geese: Spring Stopovers Are the Weak Links in Meeting Targets for Breeding. J Ornithol 148:501–514.

Ferreras, P (2001). Landscape Structure and Asymmetrical Inter-Patch Connectivity in a Metapopulation of the Endangered Iberian Lynx. Biological Conservation 100:125–136.

Fortin, D, Beyer, HL, Boyce, MS, et al. (2005). Wolves Influence Elk Movements: Behaviour Shapes a Trophic Cascade in Yellowstone National Park. Ecology 86:1320–1330.

Gaidet, N, Cappelle, J, Takekawa, JY, et al. (2010). Potential Spread of Highly Pathogenic Avian Influenza H5N1 by Wildfowl: Dispersal Ranges and Rates Determined from Large-Scale Satellite Telemetry. J Appl Ecol 47:1147–1157.

Garriga, J, Palmer, JRB, Oltra, A, et al. (2016). Expectation-Maximization Binary Clustering for Behavioural Annotation. PLOS ONE 11:e0151984.

Gehrold, A, Bauer, HG, Fiedler, W, et al. (2014). Great Flexibility in Autumn Movement Patterns of European Gadwalls *Anas Strepera*. J Avian Biol 45:131–139.

Graham, CH (2001). Factors Influencing Movement Patterns of Keel-Billed Toucans in a Fragmented Tropical Landscape in Southern Mexico. Conservation Biology 15:1789–1798.

Green, M, Alerstam, T, Clausen, P, et al. (2002). Dark-Bellied Brent Geese *Branta Bernicla Bernicla*, as Recorded by Satellite Telemetry, Do Not Minimize Flight Distance during Spring Migration. Ibis 144:106–121.

Guo-Gang, Z, Dong-Ping, L, Yun-Qiu, H, et al. (2011). Migration Routes and Stop-Over Sites Determined with Satellite Tracking of Bar-Headed Geese *Anser Indicus* Breeding at Qinghai Lake, China. Waterbirds 34:112–116.

Hawkes, LA, Balachandran, S, Batbayar, N, et al. (2013). The Paradox of Extreme High-Altitude Migration in Bar-Headed Geese *Anser Indicus*. Proc R Soc B 280:20122114.

Hawkes, LA, Balachandran, S, Batbayar, N, et al. (2011). The Trans-Himalayan Flights of Bar-Headed Geese (*Anser Indicus*). PNAS 108:9516–9519.

Hussey, NE, Kessel, ST, Aarestrup, K, et al. (2015). Aquatic Animal Telemetry: A Panoramic Window into the Underwater World. Science 348:1255642.

Kalra, M, Kumar, S, Rahmani, AR, et al. (2011). Satellite Tracking of Bar-Headed Geese *Anser Indicus* Wintering in Uttar Pradesh, India. J Bombay Nat Hist Soc 108:79.

Kays, R, Crofoot, MC, Jetz, W, et al. (2015). Terrestrial Animal Tracking as an Eye on Life and Planet. Science 348:aaa2478.

Keeley, ATH, Beier, P, Keeley, BW, et al. (2017). Habitat suitability is a poor proxy for landscape connectivity during dispersal and mating movements. Landscape and Urban Planning 161:90–102.

LaPoint, S, Gallery, P, Wikelski, M, et al. (2013). Animal Behavior, Cost-Based Corridor Models, and Real Corridors. Landsc Ecol 28:1615–1630.

McRae, BH, Dickson, BG, Keitt, TH, et al. (2008). Using Circuit Theory to Model Connectivity in Ecology, Evolution, and Conservation. Ecology 89:2712–2724.

Michelot, T, Langrock, R, Bestley, S, et al. (2017). Estimation and simulation of foraging trips in land-based marine predators. Ecology 98:1932–1944.

Morales, JM, Haydon, DT, Frair, J, et al. (2004). Extracting More Out of Relocation Data: Building Movement Models as Mixtures of Random Walks. Ecology 85:2436–2445.

Nathan, R, Getz, WM, Revilla, E, et al. (2008). A Movement Ecology Paradigm for Unifying Organismal Movement Research. PNAS 105:19052–19059.

Newman, SH, Hill, NJ, Spragens, KA, et al. (2012). Eco-Virological Approach for Assessing the Role of Wild Birds in the Spread of Avian Influenza H5N1 along the Central Asian Flyway. PLOS ONE 7:e30636.

Patterson, TA, Thomas, L, Wilcox, C, et al. (2008). State–space Models of Individual Animal Movement. Trends Ecol Evol 23:87–94.

Péron, G, Fleming, CH, de Paula, RC, et al. (2017). Periodic continuous-time movement models uncover behavioral changes of wild canids along anthropization gradients. Ecological Monographs 87:442–456.

Prosser, DJ, Cui, P, Takekawa, JY, et al. (2011). Wild Bird Migration across the Qinghai-Tibetan Plateau: A Transmission Route for Highly Pathogenic H5N1. PLOS ONE 6:e17622.

Richard, Y and Armstrong, DP (2010). Cost Distance Modelling of Landscape Connectivity and Gap-Crossing Ability Using Radio-Tracking Data. J Appl Ecol 47:603–610.

Row, JR, Blouin-Demers, G, and Lougheed, SC (2010). Habitat Distribution Influences Dispersal and Fine-Scale Genetic Population Structure of Eastern Foxsnakes (*Mintonius Gloydi*) across a Fragmented Landscape. Mol Ecol 19:5157–5171.

Squires, JR, DeCesare, NJ, Olson, LE, et al. (2013). Combining Resource Selection and Movement Behavior to Predict Corridors for Canada Lynx at Their Southern Range Periphery. Biol Conserv 157:187–195.

Takekawa, JY, Heath, SR, Douglas, DC, et al. (2009). Geographic Variation in Bar-Headed Geese *Anser Indicus*: Connectivity of Wintering Areas and Breeding Grounds across a Broad Front. Wildfowl 59:102–125.

Technitis, G, Weibel, R, Kranstauber, B, et al. (in preparation). On Old Tracks to New Insight: Random Trajectories on Recorded Paths.

Technitis, G, Weibel, R, Kranstauber, B, et al. (2016). An Algorithm for Empirically Informed Random Trajectory Generation between Two Endpoints. GIScience 2016: Ninth International Conference on Geographic Information Science.online.

Thurfjell, H, Ciuti, S, and Boyce, MS (2014). Applications of Step-Selection Functions in Ecology and Conservation. Movement Ecol 2:4.

Turner, GG, Reeder, D, and Coleman, JT (2011). A Five-Year Assessment of Mortality and Geographic Spread of White-Nose Syndrome in North American Bats, with a Look at the Future. Update of White-Nose Syndrome in Bats. Bat Research News 52:13–27.

van Toor, ML, Newman, SH, Takekawa, JY, et al. (2016). Temporal Segmentation of Animal Trajectories Informed by Habitat Use. Ecosphere 7:e01498.

Zeigler, SL and Fagan, WF (2014). Transient Windows for Connectivity in a Changing World. Movement Ecol 2:1.

Zeller, KA, McGarigal, K, Cushman, SA, et al. (2014). Sensitivity of landscape resistance estimates based on point selection functions to scale and behavioral state: pumas as a case study. Landscape Ecology 29:541–557.

Zeller, KA, McGarigal, K, Cushman, SA, et al. (2016). Using step and path selection functions for estimating resistance to movement: pumas as a case study. Landscape Ecology 31:1319–1335.

